# Why they occur here but not there – a case study of Great Spotted Woodpeckers in the city of Munich

**DOI:** 10.1101/2025.07.06.663361

**Authors:** Wolfgang W. Weisser, Andrew Fairbairn, Maximilian Mühlbauer, Lea Guthmann, Annika Arndt, Yannick Bauer, Livia Caravitis, Felix Fischer, Virginia Herre, Johanna Kaucher, Karen Klein, Isabel Kleinschroth, Sandra Niederlechner, Inga Peters, Leona Schwering, Amelie Zeller, Sebastian T. Meyer

## Abstract

Many animal species can occur in the urban environment. The factors determining whether a particular species can live in a particular place in the city are, however, often not known. Such knowledge is needed to be able to design a built environment that is suitable for species.
The Great Spotted Woodpecker, *Dendrocopos major*, breeds in self-made cavities in trees and regularly occurs in cities. Here, we used a number of approaches to elucidate the factors that influence the occurrence of woodpeckers in the build-up area of the city.
We studied the occurrence of *D. major* on 103 squares in Munich, Germany and linked their presence to a number of square features such as square size, position in the city, tree occurrences, and a number of other variables. We also observed the presence of woodpecker cavities and undertook behavioural observations to study tree use.
Woodpeckers were observed on 60% of the squares. A random forest model showed that the number of humans on squares negatively affected woodpecker occurrence, while a number of tree-related variables, such as tree density, tree diversity, the number of old trees (>60 cm DBH), and the sum of DBH of trees on the square, positively affected occurrence. Some other variables were also important, such as the amount of green in the surroundings and the distance to the city centre.
Nests were found only in trees with a DBH>20 cm, and there was a distinct preference for larger trees, whereby a range of species was accepted. Behavioural observations showed that woodpeckers used a wide variety of trees, including small ones. Particular species and trees with a DBH>20 cm are preferred.
Our study shows that humans can design public places for the presence of the Great Spotted Woodpecker by providing a locally high abundance of trees of different species with total DBH close to 25m. Trees in the surroundings and the presence of softwood trees with >40 cm will make it more likely that woodpeckers will use such sites in the city.

**Statement on inclusion:** Our study brings both established scientists with permanent faculty positions with Ph.D.-students and students of the 2^nd^ year of their B.Sc.-studies or 1^st^ or 2^nd^ year of their M.Sc.-studies for whom this work is the first experience in the publication of scientific results. Students were particularly encouraged to contribute to question and hypothesis development as well as analysis and interpretation.

## Introduction

In recent years, there has been an increasing interest in the occurrence of animal species in the urban environment (Rega-Brodsky *et al*. 2022). Today, it is clear that many species can live within city boundaries (Sweet *et al*. 2022). However, what has also become clear is that a city is not a homogeneous entity and that the distribution of different urban elements plays an important role in where a particular species can be found. For example, the seminal meta-analysis by Beninde, Veith and Hochkirch (2015) found that green ‘patch’ area and the presence of corridors were most important to explain patch species richness for a number of taxa. More recent studies have thus started comparing different land uses within the cities, or focused on particular habitats such as gardens, but also public squares (Fontaine *et al*. 2016; Melliger, Rusterholz & Baur 2017; Baldock *et al*. 2019; Korányi *et al*. 2021; Mühlbauer *et al*. 2021; Rega-Brodsky *et al*. 2022; Fairbairn *et al*. 2024). These studies have found that various local features, such as overall greenness, total vegetation, vegetation structure, or human management, but also the presence of potential habitats in the surroundings, affect alpha diversity of a focal patch (Melliger, Rusterholz & Baur 2017; Baldock *et al*. 2019; Korányi *et al*. 2021; Mühlbauer *et al*. 2021). Responses to these features often differed among the different taxa considered, not only between plants and animals, but also among animal groups such as grasshoppers and butterflies (e.g., Melliger, Rusterholz & Baur 2017). Because animals are an important part of urban nature and generally liked by people (Sweet *et al*. 2023), the knowledge of why a particular animal species can live at a specific place in the city can be used to design the urban environment in such a way that the species can occur there, not only in parks, but also in the built-up area of the city (Weisser & Hauck 2024). In this paper, we analyse why the Great Spotted Woodpecker, *Dendrocopos major* L., a species that is fairly common in the urban environment, occurs in some sites in the city, but not in others. Specifically, we concentrate on public squares, which are urban elements designed with human use in mind but that can also host various animal species (Fairbairn *et al*. 2024).

Woodpeckers are part of the family *Picidae* that usually nest and roost in tree holes that they excavate themselves. Woodpeckers have evolved in forests and are generally considered forest specialists. However, many woodpecker species can be found in fragmented and disturbed urban and suburban landscapes (Kotaka & Matsuoka 2002; Morrison & Chapman 2005; Kajtoch & Figarski 2017; Bovyn *et al*. 2019; Catalina-Allueva & Martín 2021). For example, in Europe, the Great Spotted Woodpecker, the Green Woodpecker *Picus viridis* L., but also other species of woodpeckers, frequently occur in cities. While continuous green space of at least 10-35 hectares has been suggested to be the minimum size necessary to support most urban bird species, urban parks and green spaces are generally much smaller and still support a number of bird species (Lepczyk *et al*. 2017). Woodpeckers can be found on patches well below 10ha, nesting in trees lining urban streets and foraging in apartment block courtyards, clearly able to find the necessary resources within the urban fabric (von Blotzheim, Bauer & Bezzel 1966). Some research suggests that urban woodpeckers may be limited by factors such as the availability of certain tree species, by tree density (Sandström, Angelstam & Mikusiński 2006), patch size (Morrison & Chapman 2005), or the availability of deadwood (Catalina-Allueva & Martín 2021). However, studies of urban woodpeckers have so far mostly focused on urban parks, cemeteries, or other larger green spaces where such resources may be more concentrated.

The Great Spotted Woodpecker is a mid-sized woodpecker that is common throughout Europe, including Germany, and regularly occurs in urban areas (von Blotzheim, Bauer & Bezzel 1966). They have a varied and seasonal diet, taking a variety of arthropods, carrion, nuts, berries and the eggs and young of other birds. In winter, cones of spruce and other conifers make up a large part of the diet, although, again, in the city, individuals are flexible in their diet (von Blotzheim, Bauer & Bezzel 1966). Individuals breed in self-made tree cavities and also spend most nights roosting in a cavity (von Blotzheim, Bauer & Bezzel 1966). In the city, abandoned holes of the woodpecker are important for many other species in this cavity-poor environment (Kotaka & Matsuoka 2002), which have to be substituted by nestboxes when appropriate trees and woodpeckers are not present. While the Great Spotted Woodpecker is one of the best-studied European woodpeckers, most information from cities again concentrates on parks and cemeteries (e.g. Pasinelli 2006; Figarski & Kajtoch 2018). For example, Schütz, Reckendorfer and Schulze (2017) found for parks in Vienna, Austria, that park size was the most important local variable affecting the occurrence of the Great Spotted Woodpecker, with woodpeckers occurring in 20 out of the 29 parks (69%) in the city.

In this study, we focus on the occurrence of the Great Spotted Woodpecker (henceforth ‘woodpecker’) on urban squares, which are a core element of cities and are designed by humans for human use (Mühlbauer *et al*. 2021; Fairbairn *et al*. 2024). We were interested in elucidating the factors that determine whether the woodpecker can occur on an urban square. Our approach was as follows: first, we used data from a bird survey conducted on 103 urban squares in Munich, Germany (Mühlbauer *et al*. 2021; Fairbairn *et al*. 2024) and extracted data on the occurrence of the woodpecker as well as square features. We additionally extracted data on vegetation on the 103 squares from square surveys that we considered relevant for the woodpecker. We then carried out additional observations of woodpecker presence and the behaviour of individuals, to further elucidate the use of trees by woodpeckers for foraging and making holes.

We asked the following questions:

1. What features of the urban squares are correlated with the presence of the woodpecker on urban squares, including both vegetation-related variables as well as other variables such as the number of humans visiting the square?
2. Is there a difference in square features between squares where woodpeckers are observed more frequently and those where they are rarely or never observed?
3. What trees are used by woodpeckers with respect to species and diameter?

## Materials and Methods

### Selection of urban squares and square features

We selected 103 squares in Munich, Germany, out of a candidate list of 354 public squares (Supplementary Material Fig. S1; Mühlbauer *et al*. 2021; Fairbairn *et al*. 2024). We used stratified random sampling while accounting for square size, the distance to the city centre and the presence of trees. We then characterised squares by a number of different features, including both vegetation-related variables and ‘grey’ characteristics relating to square design (Supplementary material, Table S1). A full description of the methodology is given in Mühlbauer *et al*. (2021) and Fairbairn *et al*. (2024). Briefly, the greenness of the squares was calculated as the proportion of a square’s area that was vegetated (Normalized Difference Vegetation Index – NDVI values ≥ 0.2). Square size, position in the city, proportion of sealed areas, number of streets bordering the square, the presence of water and night-time illumination (Artificial Light at Night – ALAN) were assessed from different digital sources. Using on-site visits, we measured the planted vegetation on each square as flower beds, grass (grass-dominated low vegetation), shrubs and trees. For trees, the species and individual tree size (DBH) were registered. Old trees (DBH ≥ 60 cm) were counted separately. Because trees are important for woodpeckers, we calculated a number of variables such as tree species richness, Shannon diversity of trees, tree (and old tree) abundance, average and total tree DBH of the square. For shrubs, we estimated volume, and for lawns and flower beds, we estimated the area in square meters. For many features, we calculated both the total area and the proportion of the square they occupied. In this analysis, we used the total area of the square as a variable and used the proportion of the different vegetation types as further variables (Table S1). Human usage of squares and the presence of domestic animals were quantified by counting estimating the number of humans on each square during each bird survey. Because connectivity and availability of habitat in the surrounding area have been shown to be important for biodiversity in urban environments, we also calculated the percentage of green area in a buffer of 1000 m around the border of each square.

### Occurrence of woodpeckers in urban squares

All 103 squares were sampled in seven sampling rounds: three sampling rounds during the breeding season (April to July 2017), one in autumn (October 2017), and three sampling rounds in winter (December 2017 to February 2018), as described in Mühlbauer *et al*. (2021). Each round lasted about 14 days. Survey time per square was standardized to 20 minutes. Predefined routes were walked with a constant pace that was adjusted so that the whole route was covered exactly once. All bird species were recorded within a 25 m buffer along the track. Here, we use the data on the occurrence of the Great Spotted Woodpecker from this study.

### Mapping of woodpecker occurrences and woodpecker holes on squares in 2020

In 2020, all squares were visited again in the summer period, and both the occurrence of woodpeckers and woodpecker holes were recorded. When a tree with a hole of the Great Spotted Woodpecker was encountered, the tree species and DBH were noted down. The cavity data were used together with the tree data used in Mühlbauer *et al*. (2021) to compare the distribution of trees with cavities to the overall number of trees of all species.

### Woodpecker use of trees in 2024

From 13.05.2024 and 03.06.2024, we observed tree use by 32 woodpecker individuals (17 males, 13 females, 2 juveniles). Because the aim was to obtain general information about woodpecker behaviour, we chose locations in parks and residential areas within Munich and in Freising, close to Munich (Table S2, supplementary material). Data from the birdwatching online portal “ornitho.de” as well as the sightings of woodpeckers on the 103 squares were used to identify potential sites. Initial observations were made in the Munich districts Bogenhausen, Isarvorstadt, Giesing and Ackermannbogen. In order to increase the number of woodpeckers observed, further places and parks were subsequently visited. The potential sites were visited and, if the woodpecker was encountered, the observation started. All individuals were observed for at least two hours (mean ± sd: 135 ± 36.9 minutes).

Observations were done in groups of three people. The first person had the task of following the woodpecker as closely as possible and observing its behaviour with binoculars. Two further people marked all the trees used by the woodpecker with road chalk and took notes. Woodpecker cavities were often the starting point for the observations. As it was not possible to map the trees in the order in which they were used, or to determine the exact duration of stay per tree due to the short residence times of woodpeckers on trees and the speed of flights, the number of visits per tree was recorded instead, and it was only noted if a tree visit was longer than two minutes. Behaviours observed were foraging, drumming, feeding young, calling, grooming, and resting/stopping. The marked trees were given a unique ID and then entered into a Google Maps project with their exact location. For all trees visited, the DBH and species were recorded. The observations were stopped when a woodpecker individual could be observed for at least two hours. In order to collect sufficient data, the observations of single individuals were sometimes spread over several days.

In order to determine tree preferences of the woodpecker in terms of tree species and DBH, an observation area was defined based on the markings of the trees in Google Maps (example in Fig. S3, supplementary material. Boundaries of the observation area were determined by connecting the outermost observation points and by making use of natural boundaries of the area, such as paths or woodland edges. We then recorded species and DBH of all trees with a DBH>5cm within the observation area for trees.

### Statistical analysis of woodpecker occurrences on the 103 squares

All analyses were conducted in R (version R-4.1.1) using RStudio (version 2021.09.0). To determine how different square features affected the presence of woodpeckers on the 103 urban squares, we used the monitoring data from 2017/18 and 2019. To test the effect of greenness (NDVI) on the occurrence of woodpeckers on squares, a logistic regression was fitted. To determine the predictive power of the different square features on the occurrence of woodpeckers on squares, we ran random forest models, which allowed us to include all selected variables in the model (cf. Fairbairn *et al*. 2024). While random forest models are relatively robust against the correlation of individual predictor variables, we nevertheless reduced these correlations by conducting a principal component analysis (PCA, using the *prcomp* function in the *stats* package) with all square features (Fig. S2, supplementary material). All absolute measures of area, except for the overall area of each square, were removed, and only proportional variables were used. From clusters of closely related variables, we selected those which we consider to be biologically most interpretable for the analysis. The following variables were thus not used in random forests: proportion of green, shrub volume, proportion of grass, abundance of trees, tree index, abundance of old trees, proportion of unsealed surface, proportion of flowers, density of tree richness (Table S1). The proportion of green, shrub volume, proportion of unsealed surfaces, proportion of sealed surface and proportion of grass were all correlated with each other and other variables. Some variables, for example, the proportion of green and unsealed surface, describe much of the same thing, and the proportion of sealed surface is directly negatively correlated with several of the variables representing vegetation properties. The proportion of flowers was removed as they are not considered biologically relevant for woodpeckers. The abundance of trees and the abundance of old trees were removed as they are size-dependent. The tree index is a combination of tree density and diversity and was removed for interpretability. The density of tree richness was removed for interpretability.

To run the random forest models, we used the *caret* package (Dennis *et al*. 2017) with increasing forests with 1001 trees. Final models were created using leave-one-out cross-validation (LOOCV). We tuned the mtry parameter (number of variables randomly sampled as candidates at each split) by evaluating a predefined set of values (2, 4, 6, 8, and 10). Variable importance was calculated using the varImp function in *caret*, which calculates the percent increase in Mean Square Error (%IncMSE). The %IncMSE score for a variable is the difference between the final model MSE and the MSE after permuting the order of the values of that predictor. Variables with higher values of %IncMSE have a larger effect on the predictive performance of a model. The variable importance was then scaled, ranging from 0 to 100. Finally, the direction of the effect (positive or negative) was determined by inspection of the partial plots.

### Characterisation of squares differing in woodpecker observations

Based on the number of times a woodpecker was encountered during a square visit in 2017/2018 (max. 5 times in the 7 visits to the square), we divided squares into ‘frequently occupied’ (4-5 times observed, 11 squares), ‘sometimes occupied’ (1-3, 51), and ‘never occupied’ (0, 41). To test for significant differences between squares where woodpeckers were frequently, sometimes, and never observed, we first conducted a Kruskal-Wallis test for each square property. To determine which specific groups differed, we conducted a pairwise Wilcoxon rank-sum test.

### Statistical analysis of woodpecker preference for trees of different species and DBH

To examine whether the presence of woodpecker cavities in trees on squares or woodpecker activity was influenced by tree species and diameter at breast height (DBH), we constructed a three-way contingency table and fitted hierarchical log-linear models using the *loglm* function from the MASS package (Venables & Ripley 2002) in R.

Tree species, DBH category (10 cm bins) and cavity presence (yes/no) or activity (yes/no) were treated as categorical variables. Tree species encountered with fewer than five individual trees were grouped into the category “rare”. Due to the sparse nature of our contingency table, we employed Monte Carlo methods to obtain more reliable p-values. Specifically, we used Monte Carlo simulation to estimate p-values for both the hierarchical log-linear models and subsequent chi-square tests of independence. For the log-linear models, we assessed model fit using Monte Carlo simulation with 10,000 replicates. After evaluating the significance of the three-way interaction, we proceeded to test pairwise associations using two-way contingency tables and Pearson’s chi-square tests with Monte Carlo simulation (10,000 replicates) to address potential violations of asymptotic assumptions in sparse tables.

Additionally, to test whether woodpeckers showed preferences for particular wood types, we classified tree species as either hardwood or softwood based on their mean wood density.

Using the Global Wood Density Database (Zanne *et al*. 2009), accessed via the BIOMASS package in R(Wagenführ & Scholz 2018). Species with a mean density above 0.55 g/cm³ were classified as hardwood. We examined the relationship between wood type and cavity presence or woodpecker activity using a two-way contingency table and Pearson’s chi-square test of independence.

In a final test to determine if woodpeckers preferentially feed in specific tree species or sizes over those used for other activities, we performed the same set of analyses as described above.

## Results

### Occurrence of woodpeckers on squares

In 2017/2018, the woodpecker was observed on 62 out of the 103 squares (60.2%) when all seven sampling rounds were considered together. In summer, woodpeckers were observed on 49 squares (47.6%, three sampling rounds), and in winter on 28 squares (27.2%, three sampling rounds). On 23 squares, woodpeckers were observed both in summer and in winter. Cavities of the woodpecker were observed on trees in 25 squares, in 80 out of the 6235 trees mapped on the squares (1.3% of the tree population).

### The influence of square features on woodpecker occurrence

A number of square features significantly predicted woodpecker occurrence on squares (Fig. 1, Supplementary material Table S3). First of all, the probability of encountering a woodpecker strongly increased with increasing greenness of the square (Fig. 1a). When the different square features (without overall greenness) were entered in the random forest model, the number of humans present on the square turned out to be a very significant predictor, both when woodpecker occurrence was based on all visits as well as when summer and winter were analysed separately. In all cases, an increasing number of humans was correlated with the decreasing probability of the woodpecker being encountered on the square. In summer, the importance of humans was slightly lower; instead, the total tree DBH and proportion of shrubs became the most important variables, positively affecting the occurrence of woodpeckers on the square. Interestingly, woodpeckers occurred on all squares with a total DBH >25m (N=17, Supplementary material Fig. S5). These variables were also important when all seasons and winter were considered.

**Figure 1:**
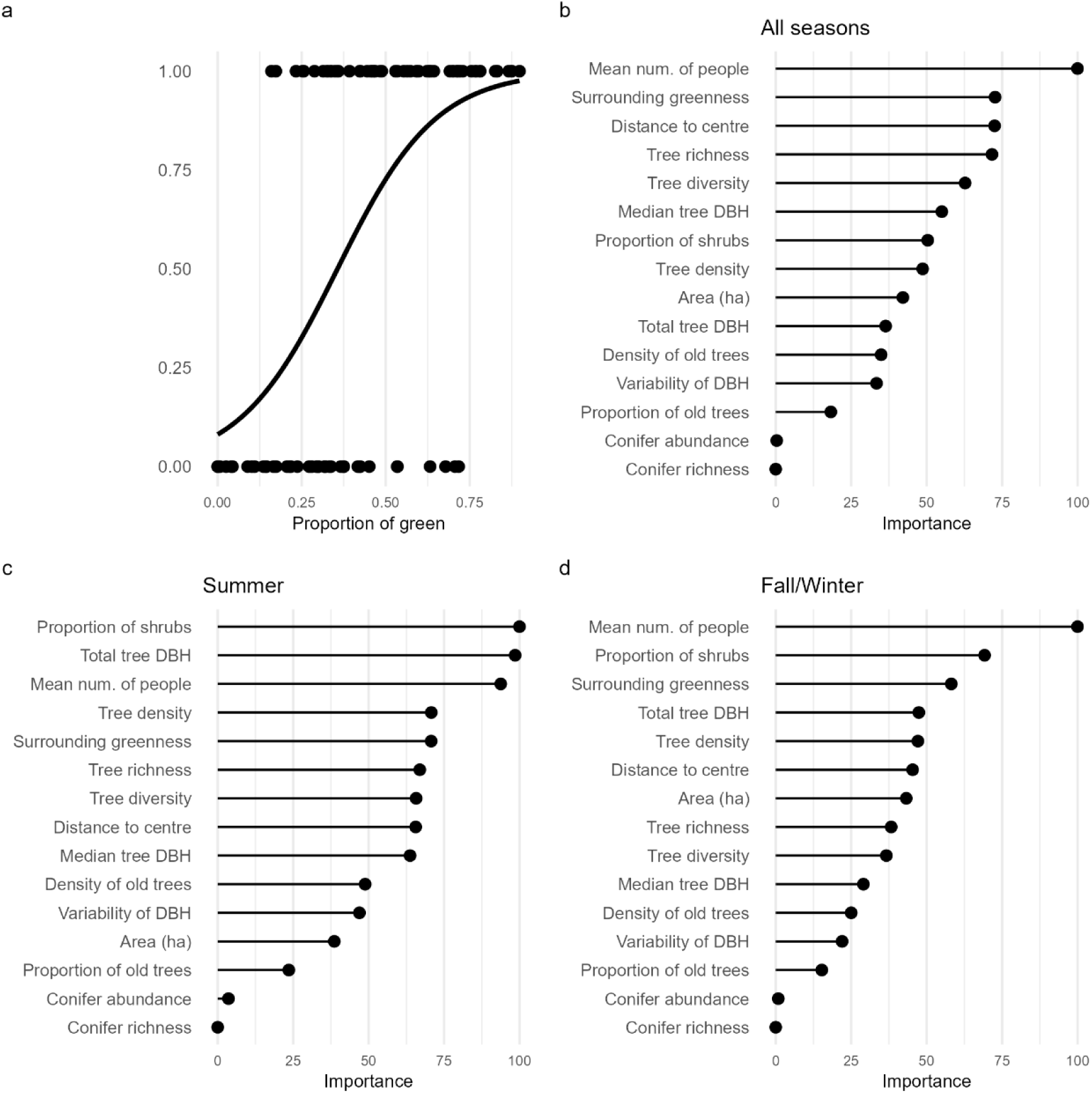
The influence of square features on the occurrence of the Great Spotted Woodpecker on squares in Munich, Germany. **a)** Effect of overall greenness of the squares (based on NDVI) on woodpecker occurrence. Woodpecker occurrence was set to 1 when woodpeckers were encountered on at least one of the seven visits to each square. The line represents a fit of a general linear model. **b)** Importance scores of different square features for the presence of woodpeckers on squares based on a random forest algorithm (7 visits). **c)** Importance scores from the random forest for woodpecker occurrence in summer 2017 (3 visits). **d)** Importance scores from the random forest for woodpecker occurrence in fall/winter 2018 (3 visits). See text for further explanation.

Tree-related variables scored high in each random forest analysis (Fig. 1). The proportion of area covered with shrubs was a significant predictor both for all samplings and the most important for summer. In addition, the distance to the city centre was also positively related to the occurrence of woodpeckers on squares, most strongly when all observations were taken together, i.e. woodpeckers were more likely to occur on the square (all other variables being equal) when the square was closer to the border of the city. The abundance and richness of conifers were not important for woodpecker presence in any season, despite the fact that woodpeckers feed on conifer cones in the winter. A likely reason is that the total number of conifer trees across all squares was only 61, corresponding to 0.01% of all trees, with a maximum of 12 trees on a square. Finally, not only were the features on the squares themselves important, but the proportion of green in a 1km radius around the square also positively affected the woodpecker presence of squares.

### Differences between squares frequently, sometimes or never used by woodpeckers

The three categories of squares (frequently occupied, sometimes occupied, never occupied) differed significantly in tree density, the proportion of green in the surroundings and the proportion of shrubs on the square (all highest for squares where woodpeckers almost always occurred, Fig. 2a-c). The three different types of squares also differed in other variables related to greenness, e.g. tree abundance or proportion of greenness (supplementary material, Fig. S6). In addition, squares where woodpeckers were never observed differed from those where woodpeckers were at least occasionally observed by being closer to the city centre (Fig. 2c), having a low density of old trees (Fig. 2d), a low tree diversity (Fig. 2e), and a higher number of humans on the square (Fig. 2g). In addition, those squares had the lowest tree richness, lowest tree index, and lowest old tree abundance (Supplementary material, Fig. S6). These comparisons emphasize that tree number, richness, and tree size, along with green in the surroundings and low disturbance from humans, are important for the woodpecker. The frequency of occurrence of woodpeckers on the squares was mirrored in the proportion of squares with tree cavities, with by far the highest proportion of squares with cavities in those squares where they were frequently observed (Fig. 2i).

**Figure 2:**
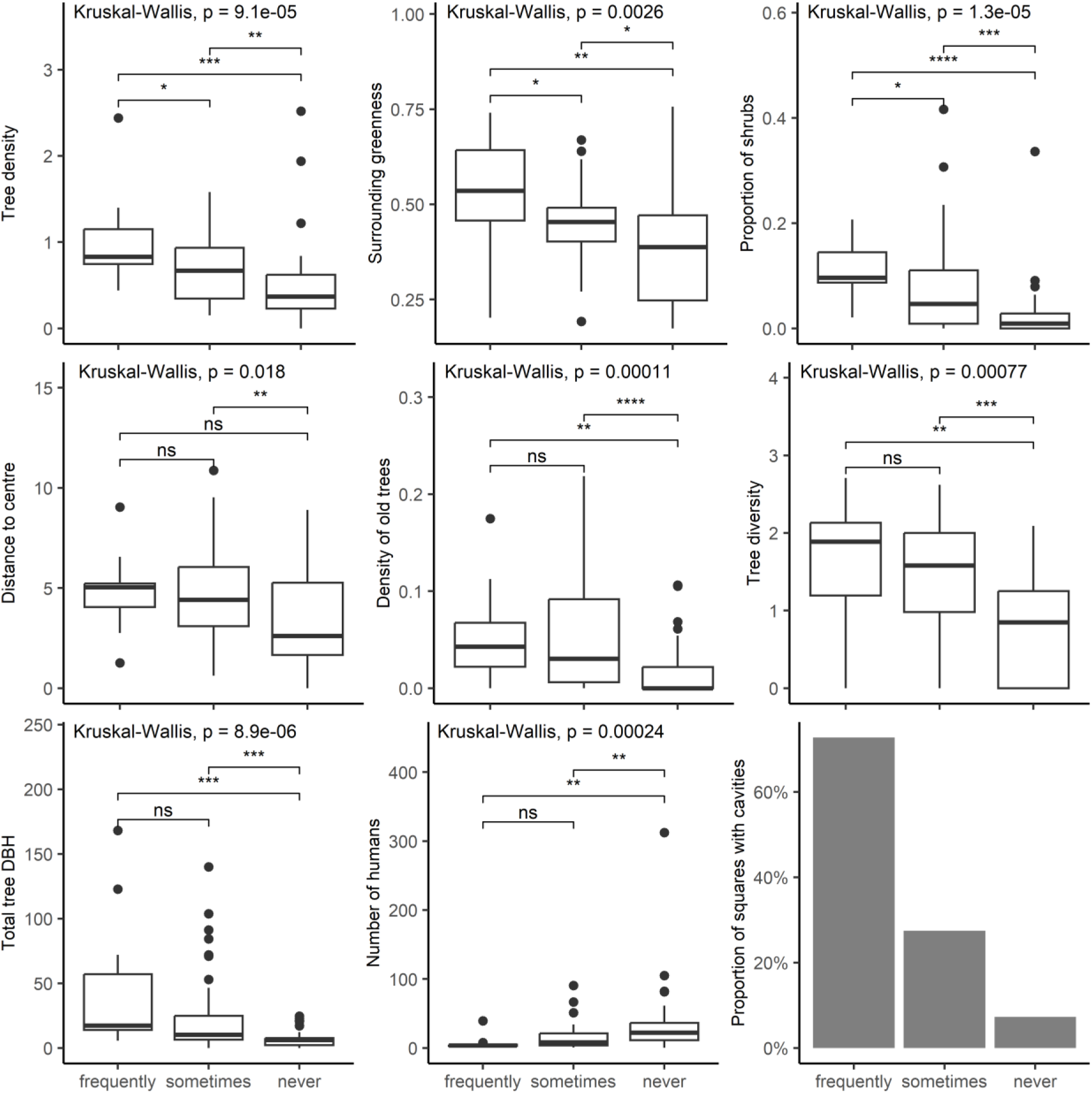
Differences in various square features between squares where woodpeckers were observed almost all of the times (at least in 4 out of 5 visits to the square, N=11 squares), where they were sometimes observed (one to three times, N=51), and where woodpeckers were never observed (N=41). Significance measures: ns: p > 0.05, *: p <= 0.05, **: p <= 0.01, ***: p <= 0.001, ****: p <= 0.0001.

### Woodpecker cavities on trees

In total, 6235 trees were counted on the squares in Munich that belonged to 78 tree species. Both *Tilia cordata* and *Acer platanoides* were present with more than 1000 individuals, followed by *T. platyphyllos* with about 600 individuals (Fig. 3a). 135 woodpecker cavities were observed on 27 squares (26%), in a total of 83 trees (range 1-7 cavities per tree).

**Figure 3:**
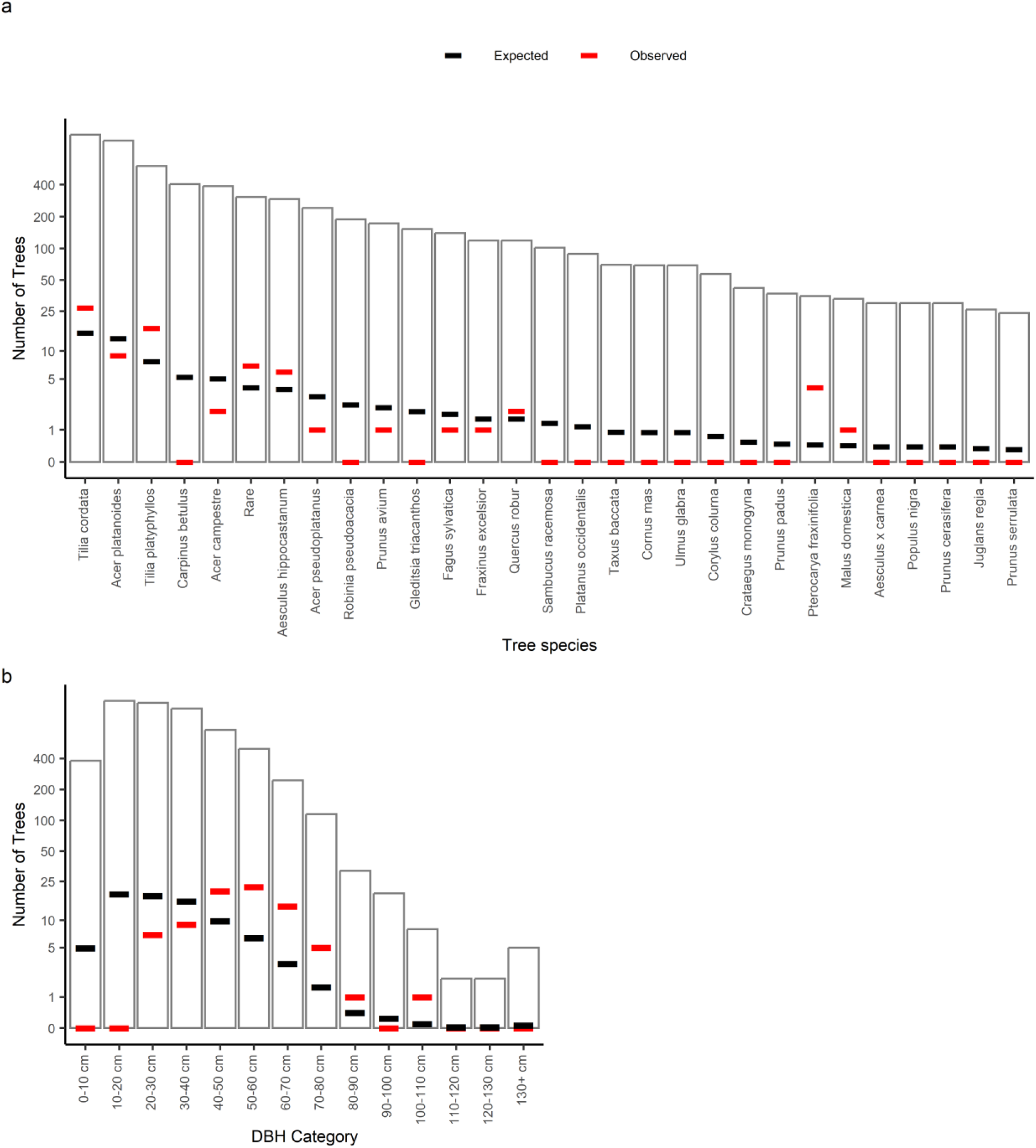
Distribution of woodpecker cavities among a) different tree species, b) trees of different diameters, on 103 squares in Munich. The black bar denotes the expected number of each diameter class/tree species based on a random distribution of cavities, and the red bar denotes the observed number. Where no red bar is visible, the value is close to zero, below the expected value and not visible.

A hierarchical log-linear analysis of cavity presence indicated that all two-way interactions between tree species, DBH category, and cavity presence were statistically significant (likelihood-ratio χ² tests, p < 0.01 for all). The presence of cavities was significantly associated with tree species (p < 0.001) and DBH category (p < 0.001). The three-way interaction among these variables was not significant, and thus, subsequent analysis focused on two-way relationships using contingency tables and chi-square tests. Expected values indicated that cavity occurrence in some species and DBH categories was much higher or lower than expected by chance (Fig. 3a, b). Thus, woodpeckers preferred trees of a certain diameter and particular tree species, but the preferred diameter did not differ significantly between tree species.

Cavities were only found on trees with a diameter of at least 20 cm. The proportion of trees with cavities was low for each diameter class (Fig. 3a). However, some rarer tree species showed a relatively high rate of cavities. *Tilia tomentosa* had a high mean DBH of 57 cm and occurred on a single square with nine individuals, where 5 (63 %) had cavities. In *Pterocarya fraxinifolia*, 10% of 39 individuals (mean DBH 52cm) had cavities, similar to *Populus tremula*. *Pinus sylvestris* was the only conifer mapped as a cavity tree; the woodpecker cavity was located in an individual with a DBH of 37 cm. In contrast, *Carpinus betulus* occurred around 400 times, without signs of woodpecker cavities. Whereas *Acer campestre* also occurred around 400 times, but had only two trees with cavities, lower than the expected five. Similarly, *Robinia pseudoacacia, Prunus avium* and *Gleditsia triacanthos* occurred at least 150 times but showed no cavities. Only a single tree with a cavity was found for *Acer pseudoplatanus, Fagus sylvatica, Fraxinus excelsior, Malus domestica, Pinus sylvestris, Populus tremula* and *Populus alba*. The number of trees on a square with a DBH>20 cm positively correlated with woodpecker presence (Supplementary material, Fig. S5).

With respect to wood type, 77 trees with cavities on squares were found in softwood, five in hardwood, and one in deadwood, while on squares there were 4479 softwood trees, 3310 of them with a DBH of 20 cm or more. There was a significant preference for softwood trees (binomial test, p<0.001). In 2024, 32 cavities were found, of which 26 were in softwood, significantly more than in hardwood tree species (binomial test, p<0.001).

### Woodpecker use of trees

In 2024, 5210 trees of 69 tree species were recorded in the observation areas. The six most common tree species were Norway maple (*Acer platanoides*) (12.6 %), hornbeam (*Carpinus betulus*) (10.4 %), field maple (*Acer campestre*) (10.3 %), sycamore (*Acer pseudoplatanus*) (9.3 %), ash (*Fraxinus excelsior*) (9.1 %) and small-leaved lime (*Tilia cordata*) (6.4 %). 37 tree species occurred less than 20 times, accounting for 3.5 % of the total number of trees. The majority of the mapped trees (94.4 %) were hardwoods, while softwoods accounted for 5.5 %. DBH ranged from 5 cm (minimum recording limit) to 170 cm (a horse chestnut, *Aesculus hippocastanum*). Average DBH was 26.6±20.6 cm, whereby 45 % of the recorded tree population had a DBH of less than 20 cm (Fig. 4).

**Figure 4:**
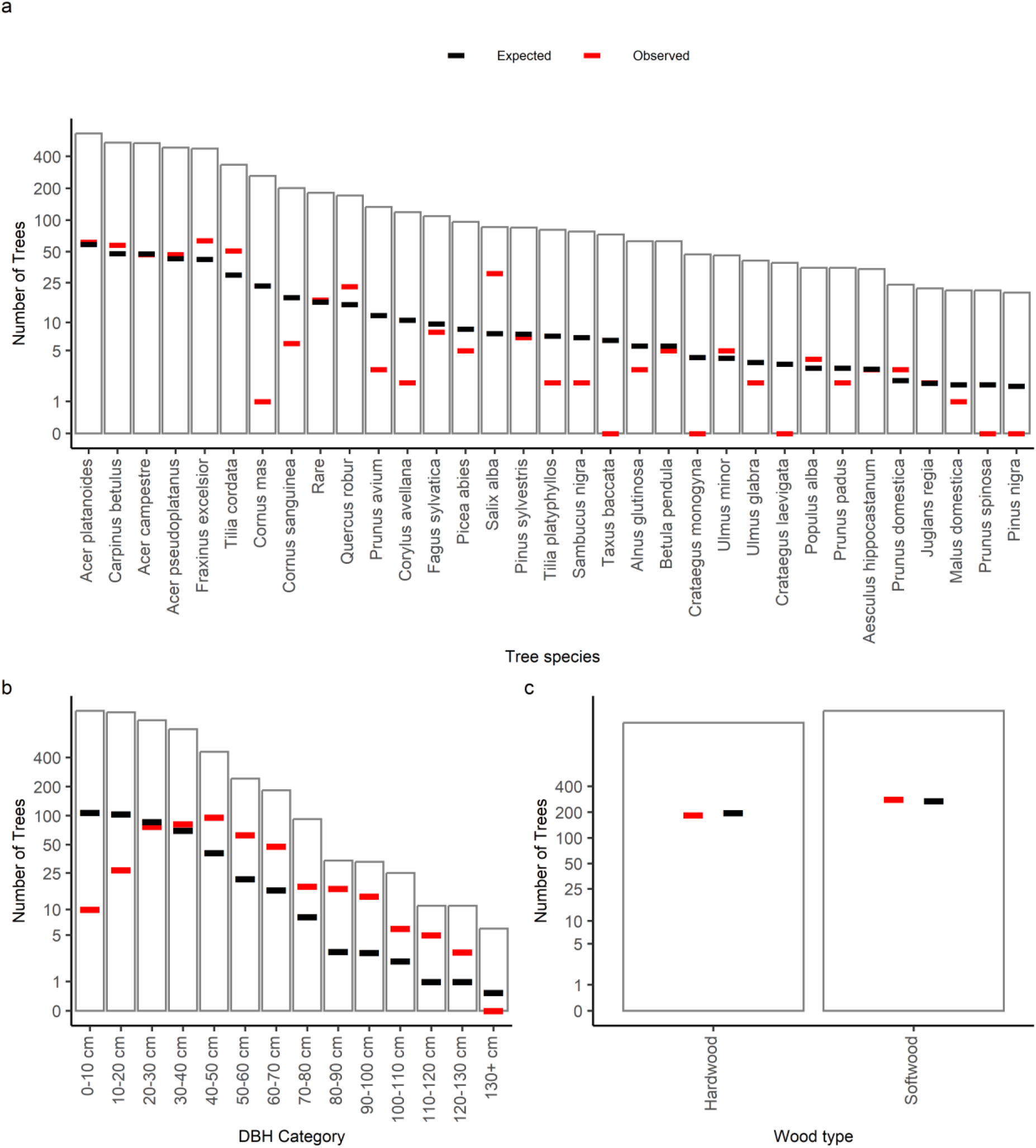
Use of trees by 32 woodpeckers in 2024 in various locations throughout Munich. Animals were observed for at least 2 hours each. During the observations, all trees where the woodpecker landed (‘used’) were marked. All trees in the observation area were identified to species and their DBH measured. a) use of different tree species, b) use of trees of different diameters c) use of trees by wood type, i.e. hardwood vs. softwood tree species. The black bar denotes the expected number of each diameter class/tree species based on a random distribution of cavities, and the red bar denotes the observed number.

During the observation periods, the woodpeckers used an average of 13±8.6 % of all trees in the surveyed areas (range 2.9-30.8%). The mean distance between the trees used and the nest trees of all woodpeckers was 45.0±35.2 m. The minimum distance between the nest and a tree that was used was 2m, and the maximum distance observed was 145m. 36 out of the 69 tree species were used by the woodpeckers.

A hierarchical log-linear analysis indicated that all two-way interactions between tree species, DBH category, and tree use were statistically significant (likelihood-ratio χ² tests, p < 0.01 for all). The three-way interaction among these variables was not significant. The chi-square tests revealed that the use of trees was significantly associated with tree species (χ² = 77.50, p < 0.001) and DBH category (χ² = 137.08, p < 0.001) (Fig. 4a, b). Thus, woodpeckers preferred higher tree DBH as well as particular tree species, but there was no interaction between tree species and DBH (Fig. 4). In contrast to the presence of cavities, woodpeckers used trees independent of whether the tree was softwood or hardwood (χ² = 1.1, p = 0.294, Fig. 4c).

The six most frequently used tree species were *Fraxinus excelsior* (64 times), *Acer platanoides* (62), *Carpinus betulus* (58), *Tilia cordata* (45), *Acer campestre* (47), and *Acer pseudoplatanus* (47, Fig. 4a). This corresponds to the most common tree species in the areas, and woodpeckers landed on those species in the same frequency as expected, or even slightly more frequently. Interestingly, one species was used much more than expected, *Salix alba*, use was four times the expected value (Fig. 4a). The shrubby species’ *Cornus mas*, *Corylus avellana* and *Cornus sanguinea*, among others, were visited less than expected, and some species such as *Taxus*, were not visited at all. In contrast to making cavities, woodpeckers also used trees <20 cm DBH, albeit at a lower than expected frequency, but all trees with a DBH of 40 cm or more (except for the rare >130 cm category) were used more frequently than expected (Fig. 4b). Overall, significantly preferred hardwood species over softwood species when accounting for tree availability (χ² = 6.33, p = 0.012; Fig. 4c).

Of 447 trees used, 223 (50.56 %) from 28 tree species were used more than once during the observation periods. Multiple visits of the same tree individual occurred most frequently in *Fraxinus excelsior*, *Acer pseudoplatanus, Acer platanoides*, *Tilia cordata, and Carpinus betulus*. The most common activity observed was foraging, followed by stopping over (using a tree to move between other trees) and calling. 367 trees of 33 tree species were used at least once for foraging. 237 trees of 31 species were used for other activities such as preening, resting, and calling. Of the trees used, we did not find that woodpeckers preferentially use specific species (χ² = 39.83, p = 0.243) or DBH categories (χ² = 17.83, p = 0.117) for foraging over other activities.

## Discussion

Many animals may occur in the city, but the reasons why they occur in one place but not in another in the urban fabric still need to be elucidated, for most species. In this study, we focused on the Great Spotted Woodpecker and linked its occurrence on public squares in Munich, Germany, to a number of variables describing the squares. In addition, we conducted behavioural observations to understand the use of trees by the woodpecker. One of our main findings was that, not surprisingly, the presence and abundance of trees are critical for the woodpecker. Our analysis emphasises the importance of a sufficient number of trees of a high enough diameter in a place, the presence of suitable tree species, as well as the presence of vegetation in the surroundings of the squares. This is consistent with recent results of (Bełcik, Woźniak & Skórka 2025), who found that the Great Spotted Woodpecker was more likely to be found in larger forest fragments in the agricultural areas north of Kracow, Poland. There were clear preferences of the woodpecker with respect to tree species and diameter at breast height, both for building a cavity and for foraging. Importantly, total DBH, i.e., the accumulated sum of all tree diameters, was a very good predictor of the occurrence of woodpeckers and of the presence of cavities on a square. However, other variables are also important, i.e. woodpeckers avoided squares close to the city centre and those with a higher number of humans using the squares. Urban planners could use these results to design public spaces where humans can observe a Great Spotted Woodpecker.

Our results are consistent with previous results showing the presence of the Great Spotted Woodpecker (Kajtoch & Figarski 2017) and other woodpecker species (Morrison & Chapman 2005) in wooded urban areas, such as parks or cemeteries. Our results go beyond these results by showing that a number of other variables, and specificities of the tree population in a site, are important in determining whether the woodpecker can be encountered in a public space. For example, if only the occurrence in summer is considered, the time when birds are breeding and feeding their young, total tree DBH, the presence of shrubs and the number of people on the square were most important for encountering woodpeckers on the square. In winter, disturbance by humans appears to be especially important. Consistent with the results of Sandström, Angelstam and Khakee (2006), we found a decreasing occurrence of the great spotted woodpeckers towards the city centre; in our study, this effect came out strongest when the entire year was considered. The Great Spotted Woodpecker uses warning calls, sometimes in a fast series (scolding) that can also be directed towards close or approaching humans (Winkler & Short 1978), but more generally, the woodpecker hides, or leaves a place in case of perceived danger (von Blotzheim, Bauer & Bezzel 1966). Thus, a high density of humans appears to represent a disturbance that can prevent woodpeckers from exploiting tree resources.

Our study unravelled clear behavioural preferences of woodpeckers towards particular tree species and tree DBH, but also showed that woodpeckers use many trees in a site, not just a few large ones. The Great Spotted Woodpecker is often described as a generalist as it does use many tree species and is omnivorous (von Blotzheim, Bauer & Bezzel 1966), and this has been reported repeatedly (e.g., Stański *et al*. 2020). In the summer, there is a preference for invertebrates collected from tree trunks, branches and also leaves of trees. We found that woodpeckers used a large number of tree species, but with clear preferences for *Acer*, *Carpinus*, *Fraxinus* and in particular *Tilia*, which were also the most common in the area, and there were also significant preferences (more visits than expected by chance) to some rarer species, such as *Salix alba*, *Populus* and *Robinia*. Stański *et al*. (2020), who studied the length of time that a tree species was visited, also found strong use of *Carpinus*, *Tilia*, *Populus*, but also *Quercus* and in particular Norway spruce, *Picea abies*, both of which were very rare in our squares. In that study, it was also found that woodpeckers visited smaller trees (10-20 cm DBH) and that woodpeckers preferred dead branches and were foraging higher up in the tree (Stański *et al*. 2020). The study by Stański *et al*. (2020) was, however, carried out in primeval oak-hornbeam-lime stands in the Białowieża, National Park in Poland where deadwood in general is much more common than in the urban environment, also in smaller Norway spruce trees. We did not record the height in which the woodpecker used a tree and whether a branch was dead, however, in our study, there was a clear preference for medium-sized trees between 30 and 50 cm. For cavities, the preference was even clearer, with cavities only in trees with a DBH>20 cm, with many cavities in very large trees (for urban standards), with a DBH>50 cm. Such large trees were used even if the tree species was rare, and softwood tree species were generally preferred. Our study thus emphasizes that in the urban environment, small trees (DBH<20 cm) are of less value to the woodpecker. Large trees are rare in urban environments, including in this study area, and their presence depends on proper tree care and the protection of existing large trees (Croeser *et al*. 2024). This again is a clear message to city planners – it is necessary to have many trees to allow the Great Spotted Woodpecker to occur in a site, but these trees must be large enough. In addition, there was a clear shortage of softwood trees species, preferred for cavity making, and of conifer trees, important as food providers in winter, on the squares. While these trees are currently more common in cemeteries and parks, such could also be planted more in the urban environment.

The fact that woodpeckers were observed on fewer squares in winter emphasises that the species is relatively mobile and that the squares included in our study were smaller than the home range of the species, especially in winter. For the Great Spotted Woodpecker, home-range size varies strongly, from 40-60ha in many forests, down to 2-10 ha (von Blotzheim, Bauer & Bezzel 1966). The low number of cavities found on squares also emphasizes that woodpeckers often use a square for foraging, but does not breed or even spend the night there. It is likely that this was the underlying reason why vegetation in the surroundings of the square was an important predictor of woodpecker presence. Even though we did not further analyse the composition of this vegetation, it will also include trees, including those where the woodpeckers could build cavities. In winter, conifer cones can play an important part in the diet of the woodpeckers, in contrast to arthropods and other seeds and fruit, which can be gathered during the summer, for example, during the time when the young have to be set (von Blotzheim, Bauer & Bezzel 1966).

## Conclusions

The Great Spotted Woodpecker occurs in the urban environment, but not everywhere – probably similar to almost all species that can principally live in the city. Understanding why they occur here but not there requires elucidating the factor that enables a species to be present at a particular urban spot. Such knowledge is necessary to be able to design an urban green space for a species (Weisser & Hauck 2024). For the Great Spotted Woodpecker our study emphasizes that the presence of nesting opportunities (larger softwood trees) and sufficient foraging opportunities (many trees that are not too small) allows this species to occur on a very urban structure – the public square –, provided there is not too much disturbance and enough green in the surroundings. This becomes less likely to occur towards the city centre; however, the variability found in our study shows that it is possible to plan for woodpecker presence. Increasing the number of conifer trees that provide food in winter, which were very rare on squares, is also likely to increase the chances of having woodpeckers on squares in winter. Because the cavities of woodpeckers can be used by other species, including insects, birds and mammals and because large trees offer habitat to many species, planning for the Great Spotted Woodpecker is likely to also benefit many other species.

## Acknowledgements

Parts of the study were carried out in the framework of the Research Training Group 2679 Urban Green Infrastructure funded by the German Research Foundation (DFG), which also funded the position of AF. We thank Pauline Riegel for her support during field work.

## Author Contributions

W.W. Weisser, S.T. Meyer. M. Mühlbauer and A. Fairbairn conceived the ideas and designed methodology, and the questions and hypotheses were discussed with all authors; Maximilian Mühlbauer, Lea Guthmann, Annika Arndt, Yannick Bauer, Livia Caravitis, Felix Fischer, Johanna Kaucher, Virginia Keller, Karen Klein, Isabel Kleinschroth, Sandra Niederlechner, Inga Peters, Pauline Riegel, Leona Schwering, Amelie Zeller collected the data; A. Fairbairn, M. Mühlbauer, L.Guthmann and W.W. Weisser led the analysis of the data. W.W.Weisser and A. Fairbairn led the writing of the manuscript. All authors contributed critically to the drafts and gave final approval for publication.

## Supporting information

### Locations of public squares studied in Munich

**Figure S1:**
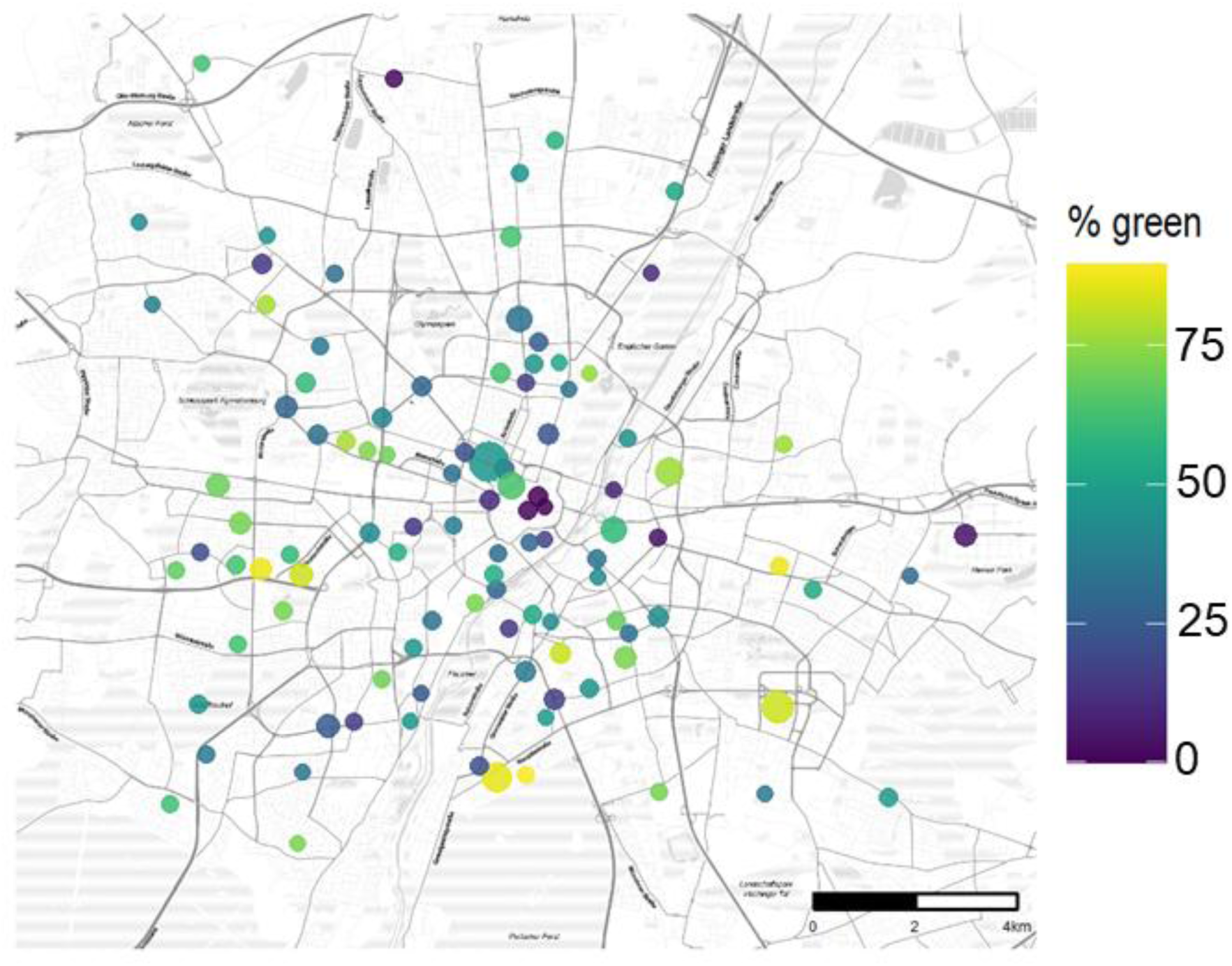
Location of the 103 squares in Munich, Germany. A detailed description of square selection is given in Fairbairn *et al*. (2024). Briefly, to delineate the borders of each square, we used aerial images and land register maps. Using the aerial images, we also calculated the Normalized Difference Vegetation Index (NDVI) for each 20 cm pixel and determined the area and proportion of greenness of each square (NDVI with a threshold ≥ 0.2). (proportion of green).

### Features measured in 103 squares

**Table S1:** Square features used in the analyses. Squares were characterized by a number of features. A detailed description of how each variable was measured is given in Mühlbauer *et al*. (2021), Mühlbauer *et al*. (2024), and Fairbairn *et al*. (2024). Table S1 lists all square features used in the analyses. The table provides the definition of each variable, range, mean ± standard deviation and indicates whether a variables was included in the random forest analysis, or excluded from further analysis due collinearity, see also the PCA-results in Fig. S2.

| Variable | Mean±sd | Range | Random forest | Description variable |
| --- | --- | --- | --- | --- |
| <u>Square size and location</u> |  |  |  |  |
| Square size (ha) | 0.92±0.98 | 0.09 - 6.71 | Included | Size of each square calculated using ArcGIS after square borders had been delineated. |
| Number of streets | 4.16±2.09 | 0 - 11 | Excluded | Number of streets connecting to the square |
| Distance from city centre (km) | 4.26±2.43 | 0 - 10.90 | Included | Distance as direct line distance from the city centre (defined as the centre of Marienplatz, the place of the townhall), to the centre of each square. |
| <u>Remotely sensed sealed, unsealed and green area</u> |  |  |  |  |
| Greenness of square | ± | 0 | | The proportion of vegetated area on a square |
| Area of green (m<sup>2</sup>) | 4633±6619 | 0 - 35657 | Included | Total area of green with NDVI ≥ 0.2 |
| Proportion of green | 0.45±0.23 | 0 – 0.9 | Included | Area of green divided by square size |
| Surrounding greenness | 0.44±0.13 | 0.174 – 0.76 | Included | NDVI with a threshold ≥ 0.2 in a buffer of 1km from the edge of each square. |
| Sealed area (m<sup>2</sup>) | 5765±4801 | 627-24094 | Excluded | 1- the unsealed area |
| Proportion area sealed | 0.7±0.2 | 0.27 – 1.00 | Excluded | Sealed area divided by square size |
| Unsealed area (m<sup>2</sup>) | 3457 ± 6205 | 0 - 42982 | Excluded | Sum of areas of lawns, shrubs, flower beds, gravel, and sand. |
| Proportion area unsealed | 0.3±0.2 | 0 – 0.73 | Excluded | The proportion of the square area that is unsealed |
| <u>Vegetation cover</u> |  |  |  |  |
| Area of grass (m<sup>2</sup>) | 2332 ± 3913 | 0 - 21464 | Included | The total area of low vegetation (except flowerbeds) on a square was assessed in a on-site visit |
| Proportion of grass | $0.21 \pm 0.16$ | 0 -0.66 | Excluded | Area of grass divided by square size |
| Area of shrubs (m<sup>2</sup>) | $725.8 \pm 1446.2$ | 0 - 8510 | Excluded | The total area of shrubs and hedges, estimated from aerial imagery on a square. |
| Shrub volume (m<sup>3</sup>) | $0.17 \pm 0.28$ | 0-1.46 | Excluded | The total volume of shrubs and hedges on a square was assessed in a on-site visit and divided by square size. Shrubs were divided in classes (<1 m, >1-2 m, >2-3 m, >3 m) and the area covered by shrubs/ hedges of each class was estimated and multiplied by height. |
| Proportion of shrubs | $0.05 \pm 0.08$ | 0 – 0.42 | Included | Area of shrubs divided by square size |
| Area of flowers (m<sup>2</sup>) | $18.1 \pm 67.6$ | 0 - 460 | Excluded | Absolute area of planted flower beds and flower tubs on a square |
| Proportion of flowers (m<sup>2</sup>) | $0 \pm 0.01$ | 0 – 0.04 | Excluded | Area of flowering plants divided by square size |
| Tree abundance | $62.7 \pm 87.3$ | 0 – 546 | Excluded | Number of trees found in the assessment of all tree individuals with DBH>7cm on the square |
| Old tree abundance | $4.3 \pm 9.7$ | 0 - 65 | Excluded | Number of trees with DBH>60cm on a square |
| Tree density | $0.64 \pm 0.47$ | 0 – 2.52 | Included | Number of trees divided by square area |
| Old tree density | $0.04 \pm 0.05$ | 0 – 0.22 | Included | Number of old trees divided by square area |
| Proportion of old trees | $0.08 \pm 0.13$ | 0 – 1 | Included | Proportion of all trees that are old |
| Tree richness | $6.92 \pm 5.96$ | 0 - 25 | Included | Derived from the assessment of all tree individuals with DBH> xxx on the square |
| Tree diversity | $1.27 \pm 0.78$ | 0 – 2.71 | Included | Shannon diversity of trees, i.e. taking abundances of tree species into account |
| Tree index | $1.59 \pm 1.5$ | 0 – 7.79 | Excluded | Index combining tree density and tree diversity |
| Median DBH (cm) | $30.3 \pm 11.1$ | 10 - 76 | Included | Median DBH of all tree individuals with DBH> 7cm on a square. When there were no trees on a square, a missing value was reported for the square (2 squares) |
| Variability of DBH | $0.46 \pm 0.19$ | 0 – 1.07 | Included | Coefficient of variation (CV) of DBH of all tree individuals with DBH> 7cm on the square (see description of DBH) |
| Sum of DBH | $19.1 \pm 29.5$ | 0 – 168.3 | Excluded | Sum of DBH of all tree individuals with DBH> 7cm on the square |
| Density of tree richness | $0.1 \pm 0.09$ | 0 – 0.24 | Excluded | Tree richness divided by square size |
| Conifer abundance |  |  | Included | The abundance of conifer trees on a square |
| Conifer richness |  |  | Included | The richness of conifer species on a square |
| <u>Other variables</u> |  |  |  |  |
| Water presence | $0.17 \pm 0.38$ | 0 - 1 | Excluded | Presence of water bodies on the square (fountains, ponds) (1 or 0) |
| Mean number of people | a) $21.49 \pm 35.9$<br>b) $14.49 \pm 43.22$<br>c) $26.89 \pm 34.89$ | a) 0.29–312.43<br>b) 0–420.67<br>c) 0–231.25 | Included | The number of humans was estimated after each bird survey as there were often too many to count them during the 20 minutes survey time. Mean numbers per season and square (a) All seasons b) Summer only c) fall & winter |

**Figure S2:**
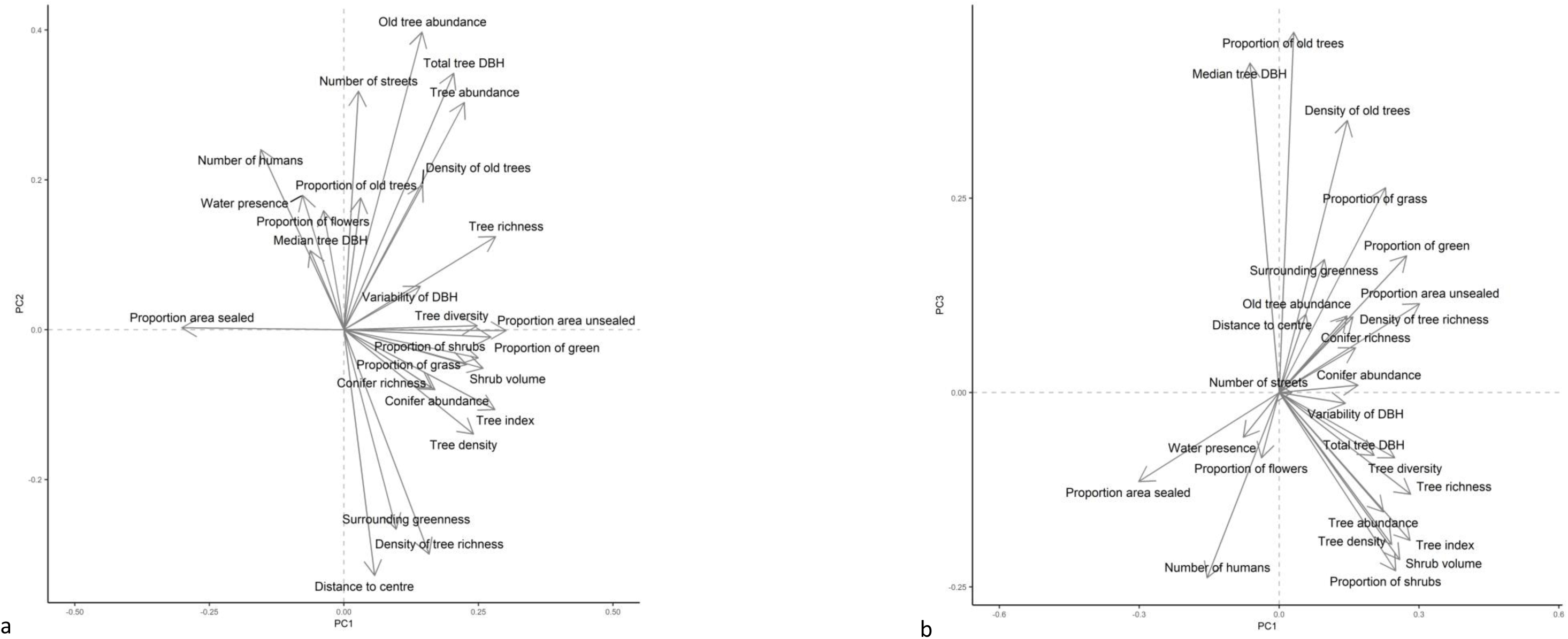
PCA of all variables. Before final section of variables for the random forest analysis, the correlation among variables was tested using principal coordinates analysis (PCA). We avoided selecting closely related variables. See text in the main manuscript for explanation. The final selection of variables is given in Table S1. a) Principal component 1 (PC1) vs. PC2, b) PC1 vs. PC3

### Behavioural observations

**Figure S3:**
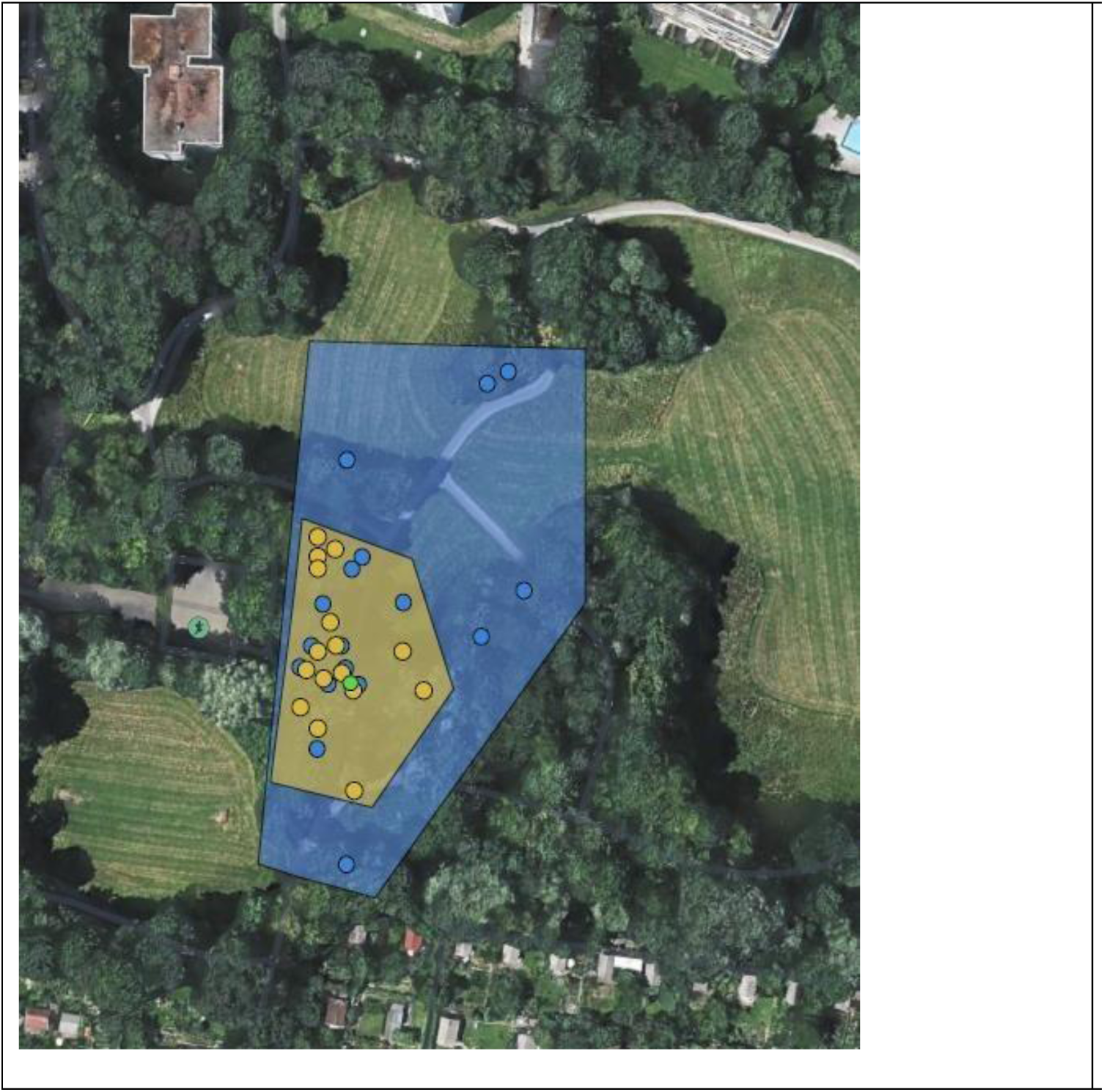
Example of observation area. Example of an observation area. In most cases, a nest of the woodpecker was at the centre of the observation area. Boundaries of an area were determined by connecting the outermost observation points and were also based on natural boundaries, such as paths or woodland edges. The dots represent the trees used (tree with nest in green, orange circles denote trees visited by a female, blue circles trees visited by a male). For all trees within the observation area with DBH > 5 cm both species and DBH were recorded.

**Table S2:** Locations of woodpecker observations. The tables shows the areas where woodpeckers were observed in 2024. Note that individuals were numbered consecutively, but some individuals could not be observed for at least 2 hours. Numbers of those individuals are therefore missing from the table.

| Individual | Sex | City/area in Munich | Location |
| --- | --- | --- | --- |
| 01 | M | Bogenhausen | Pachmayerplatz |
| 05 | M | Bogenhausen | Denninger_Anger |
| 06 | W | Bogenhausen | Denninger_Anger |
| 07 | M | Ramersdorf-Perlach | Ostpark Eisstadion NW |
| 08 | W | Ramersdorf-Perlach | Ostpark Eisstadion NW |
| 78 | J | Ramersdorf-Perlach | Ostpark Eisstadion NW |
| 09 | M | Ramersdorf-Perlach | Parkplatz am Ostpark |
| 10 | W | Ramersdorf-Perlach | Parkplatz am Ostpark |
| 41 | M | Ramersdorf-Perlach | Ostparksee |
| 42 | W | Ramersdorf-Perlach | Ostparksee |
| 44 | W | Untergiesing-Harlaching | Schrafnagelberg |
| 46 | M | Untergiesing-Harlaching | Flaucher, Osramspielplatz |
| 47 | M | Bogenhausen | Zamilapark |
| 11 | W | Isarvorstadt | Südfriedhof |
| 12 | M | Isarvorstadt | Südfriedhof |
| 13 | W | Isarvorstadt | Südfriedhof |
| 14 | M | Isarvorstadt | Südfriedhof |
| 19 | W | Lehel | Praterinsel |
| 20 | M | Lehel | Praterinsel |
| 51 | W | Untergiesing-Harlaching | Flaucher |
| 52 | M | Untergiesing-Harlaching | Flaucher |
| 53 | W | Freising | Isarauen |
| 54 | M | Freising | Isarauen |
| 21 | M | Schwabing-West | Luitpoldpark |
| 23 | W | Schwabing-West | Luitpoldpark_Borschtallee |
| 24 | M | Schwabing-West | Luitpoldpark_Borschtallee |
| 25 | M | Schwabing-West | Ackermannbogen |
| 26 | M | Milbertshofen-Am Hart | Aussiger Platz Süd |
| 27 | J | Milbertshofen-Am Hart | Aussiger Platz Süd |
| 28 | W | Milbertshofen-Am Hart | Aussiger Platz Süd |
| 29 | W | Milbertshofen-Am Hart | Aussiger Platz Nord |
| 30 | M | Milbertshofen-Am Hart | Aussiger Platz Nord |

### Results from random forest analysis of woodpecker occurrence on squares

**Table S3:** Performance metrics for random forest models. Performance metrics for random forest models predicting woodpecker presence across different seasons. All metrics are derived from leave-one-out cross-validation (LOOCV). Sensitivity represents the model’s ability to correctly identify woodpecker presence, while specificity measures correct identification of absence. Kappa values account for agreement beyond chance, with values above 0.4 indicating moderate agreement.

| Metric | All seasons | Summer only | Fall/winter only |
| --- | --- | --- | --- |
| Accuracy | 0.813 | 0.714 | 0.723 |
| (95% CI) | (0.718, 0.887) | (0.61, 9.804) | (0.622, 0.811) |
| Sensitivity | 0.900 | 0.638 | 0.707 |
| Specificity | 0.645 | 0.796 | 0.736 |
| Balanced accuracy | 0.773 | 0.717 | 0.722 |
| Kappa | 0.567 | 0.431 | 0.441 |

**Figure S4:**
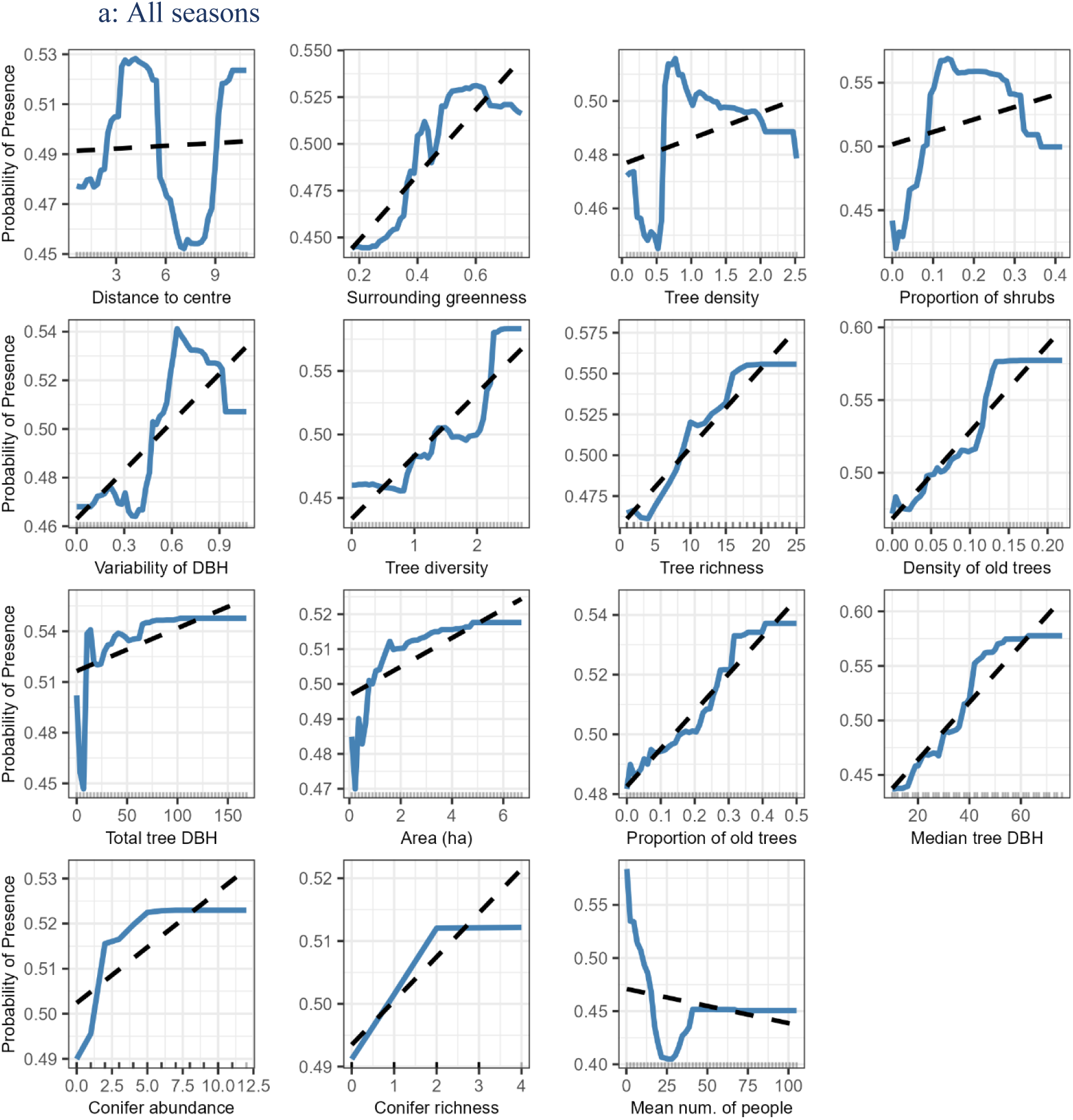
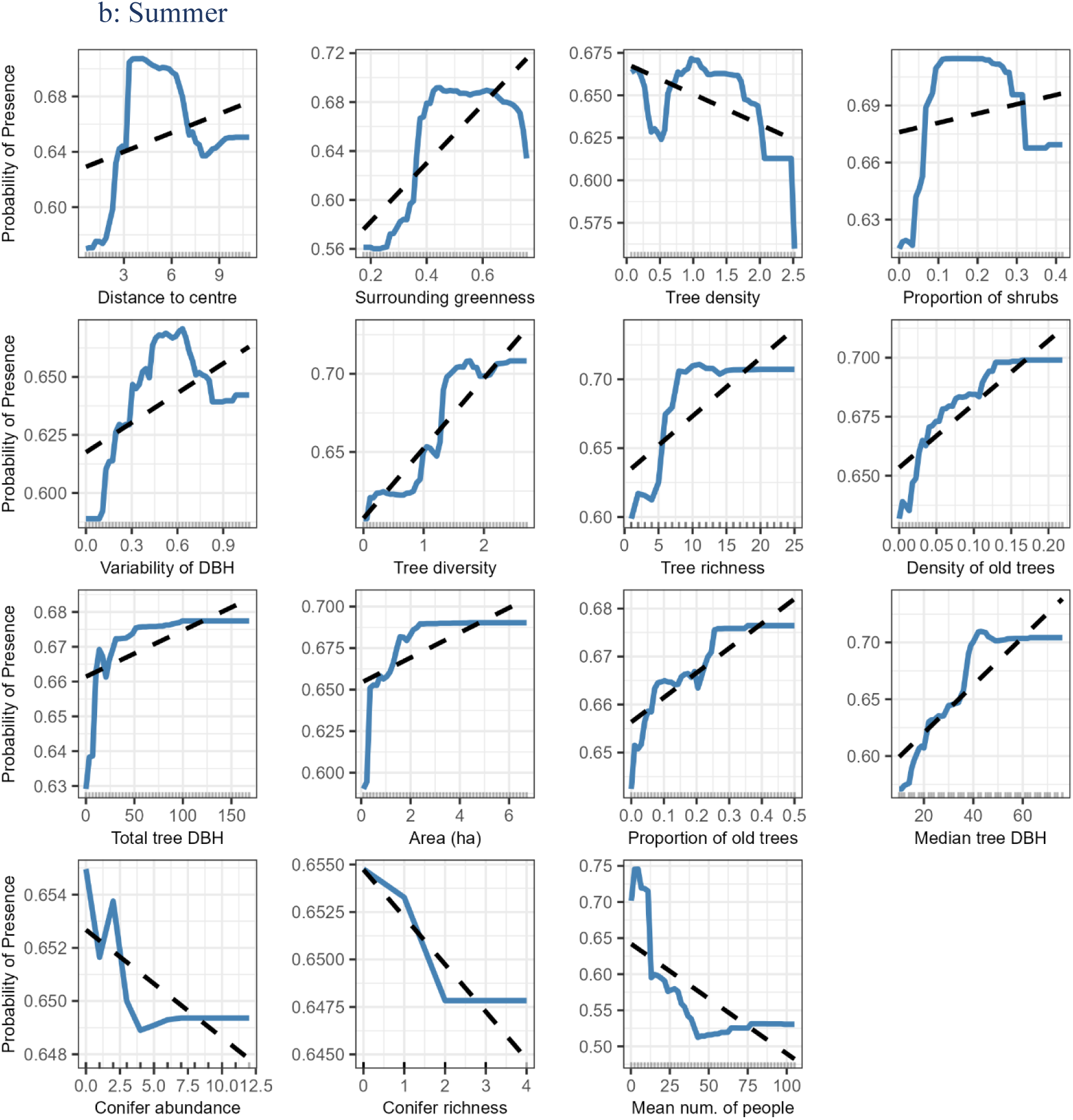
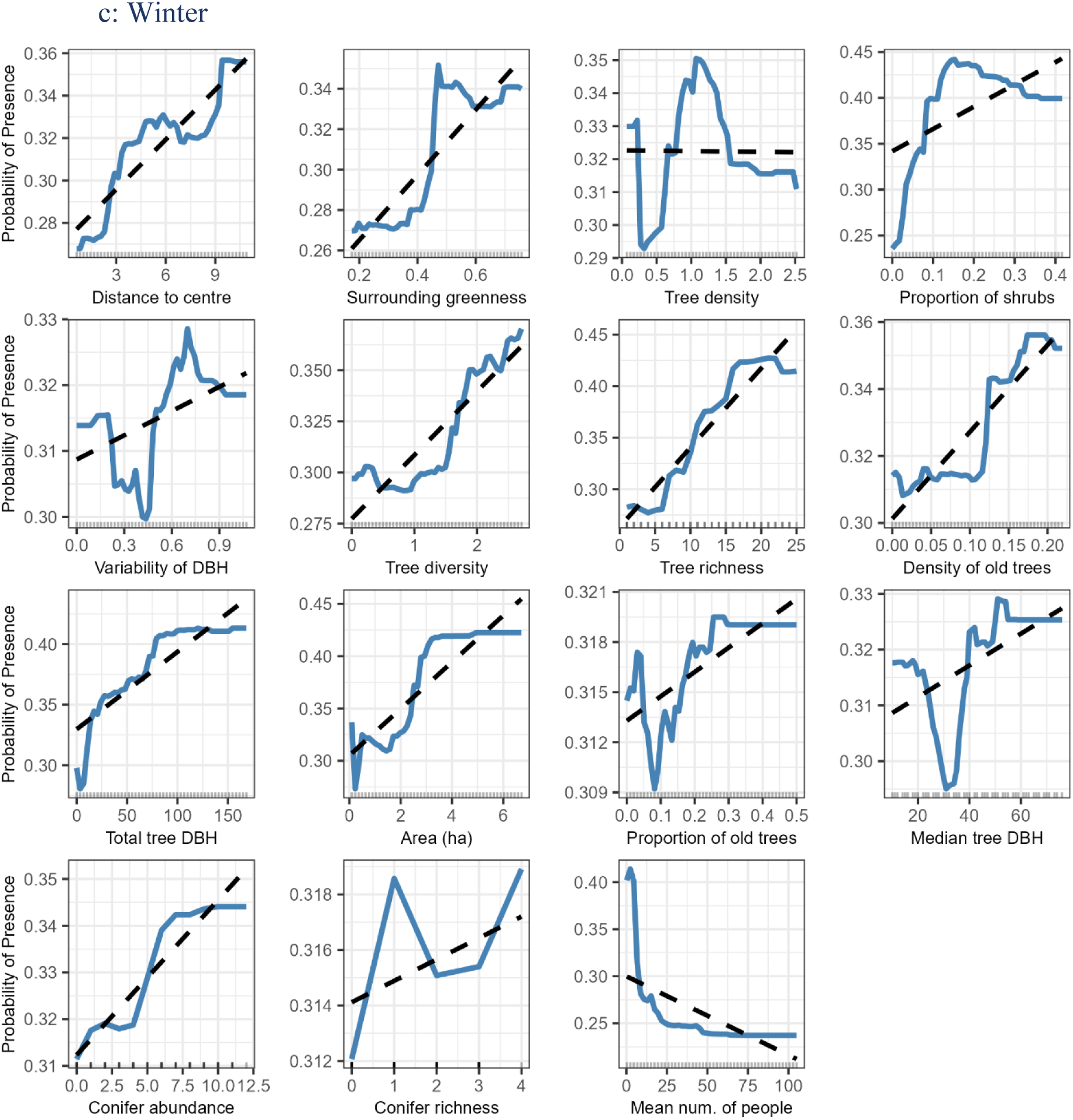
Partial dependence plots (PDPs) showing the marginal effect of each predictor variable on the probability of woodpecker presence across urban squares. Each plot displays the modeled relationship from a random forest classifier, with the blue line representing the partial dependence and the dashed black line showing a simple linear trend to illustrate the overall direction of effect. Rug marks along the x-axis indicate the distribution of observed values in the dataset.

**Figure S5:**
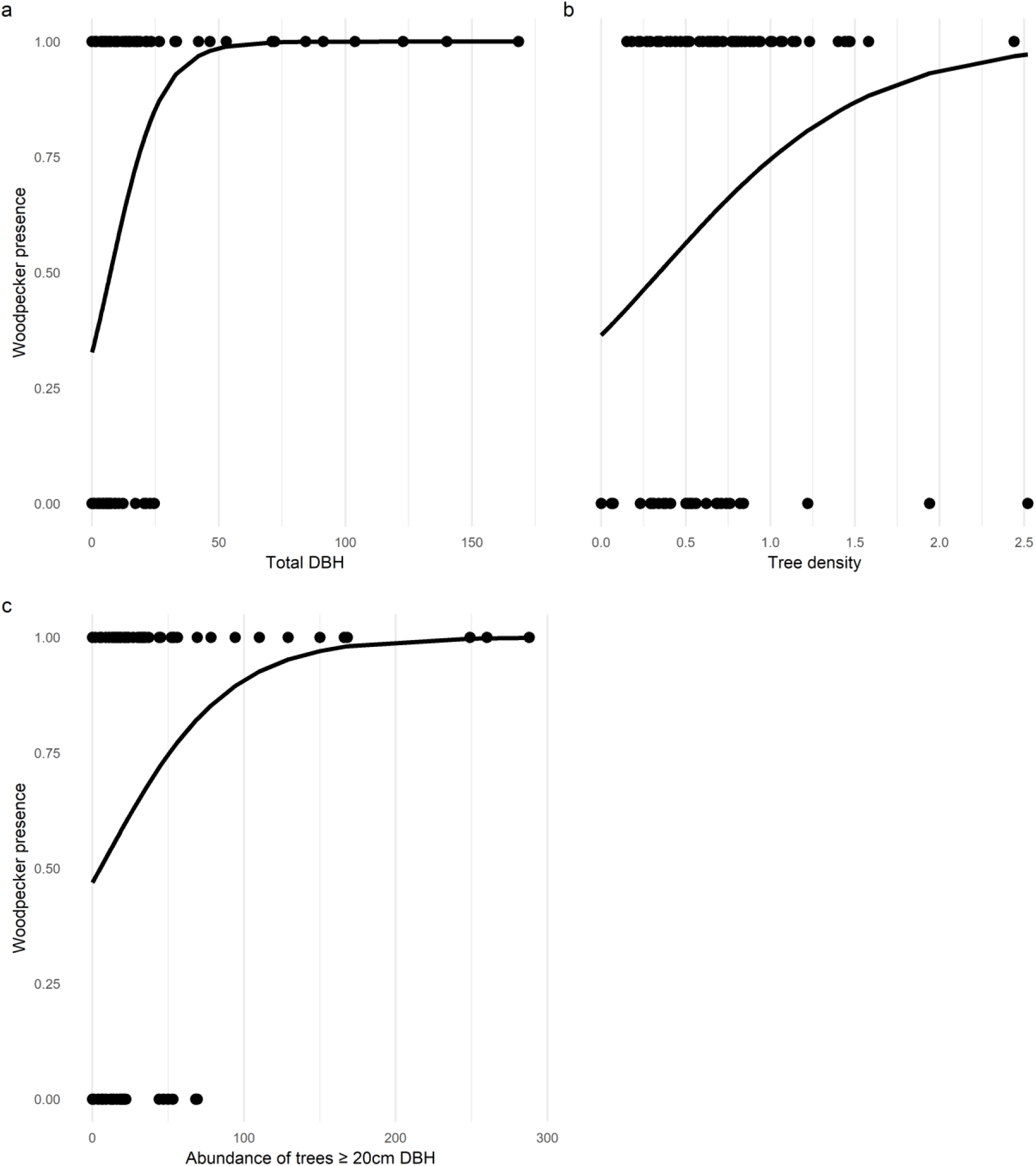
Occurrence of woodpeckers on squares as a function of total tree DBH, tree density and large tree abundance on a square. Woodpecker presence on a square as a function of a) sum of the DBH of all trees on a square (total DBH, z = 3.168, p = 0.0015), b) overall tree density (all measured trees, i.e. DBH>7cm, z = 2.779, p = 0.0055), c) abundance of trees >20cm (z = 2.479, p = 0.0132). Dots show raw data and the line is the result from a logistic regression of the respective variable on woodpecker presence on the square.

### Tree species occurring on 103 squares in Munich

**Table S4:** Total number of individuals of each tree species recorded on 103 public urban squares in Munich, Germany.

| Tree species | Number of trees |
| --- | --- |
| *Tilia cordata* | 1179 |
| *Acer platanoides* | 1038 |
| *Tilia platyphyllos* | 599 |
| *Carpinus betulus* | 405 |
| *Acer campestre* | 389 |
| *Aesculus hippocastanum* | 294 |
| *Acer pseudoplatanus* | 241 |
| *Robinia pseudoacacia* | 189 |
| *Prunus avium* | 173 |
| *Gleditsia triacanthos* | 153 |
| *Fagus sylvatica* | 140 |
| *Fraxinus excelsior* | 119 |
| *Quercus robur* | 119 |
| *Sambucus racemosa* | 102 |
| *Platanus occidentalis* | 89 |
| *Taxus baccata* | 70 |
| *Cornus mas* | 69 |
| *Ulmus glabra* | 69 |
| *Corylus colurna* | 57 |
| *Crataegus monogyna* | 42 |
| *Prunus padus* | 37 |
| *Pterocarya fraxinifolia* | 35 |
| *Malus domestica* | 33 |
| *Aesculus x carnea* | 30 |
| *Populus nigra* | 30 |
| *Prunus cerasifera* | 30 |
| *Juglans regia* | 26 |
| *Prunus serrulata* | 24 |
| *Picea abies* | 19 |
| *Pinus sylvestris* | 19 |
| *Prunus mahaleb* | 19 |
| *Betula pendula* | 18 |
| *Sorbus intermedia* | 18 |
| *Acer rubrum* | 16 |
| *Ostrya carpinifolia* | 16 |
| *Crataegus laevigata* | 13 |
| *Larix decidua* | 11 |
| *Populus tremula* | 10 |
| *Pyrus communis* | 10 |
| Ulmus minor | 10 |
| Unbekannt | 10 |
| Populus canadensis | 9 |
| Sorbus aucuparia | 9 |
| Gingko biloba | 8 |
| Tilia tomentosa | 8 |
| Acer negundo | 7 |
| Acer palmatum | 7 |
| Prunus spinosa | 6 |
| Acer buergerianum | 5 |
| Populus alba | 5 |
| Ulmus laevis | 5 |
| Abies alba | 4 |
| Quercus frainetto | 4 |
| Syringa | 4 |
| Magnolia | 3 |
| Mespilus germanica | 3 |
| Pinus nigra | 3 |
| Prunus cerasus | 3 |
| Prunus spec. | 3 |
| Salix integra | 3 |
| Ailanthus altissima | 2 |
| Larix kaempferi | 2 |
| Prunus serotina | 2 |
| Rhus typhina | 2 |
| Salix caprea | 2 |
| Sorbus torminalis | 2 |
| Tsuga canadensis | 2 |
| Abies nordmannia | 1 |
| Celtis australis | 1 |
| Ilex aquifolium | 1 |
| Rhododendron | 1 |
| Viburnum opulus | 1 |

### Differences between squares with different woodpecker occurrences

**Figure S6:**
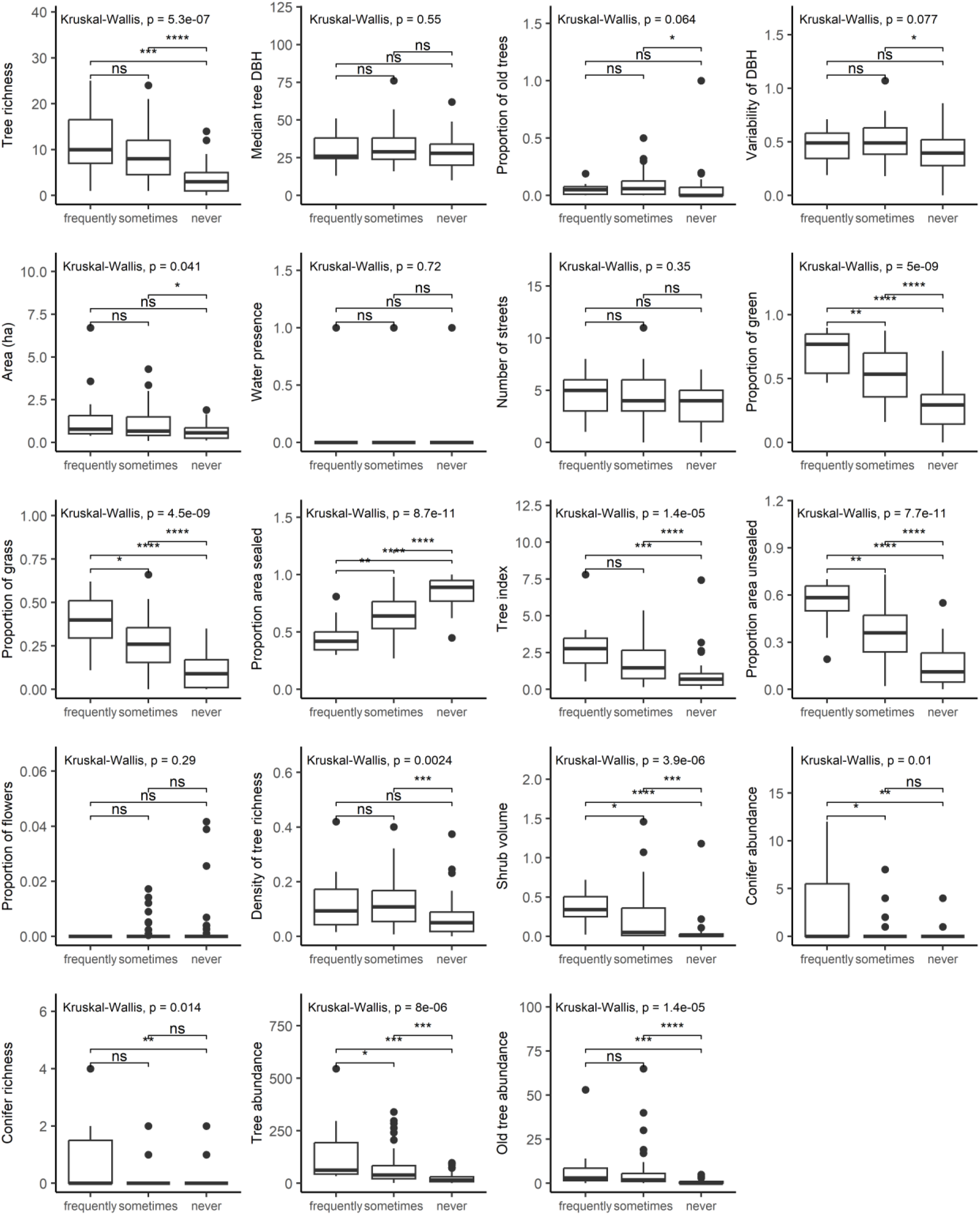
Differences in square features between squares where woodpeckers were observed in different frequencies. Differences in various square features between squares where woodpeckers were observed almost all of the times (at least in 4 out of 5 visits to the square, N=11 squares), where they were sometimes observed (one to three times, N=51), and where woodpeckers were never observed (N=41). Significance measures: ns: p > 0.05, *: p <= 0.05, **: p <= 0.01, ***: p <= 0.001, ****: p <= 0.0001.

## References

Baldock, K.C.R., Goddard, M.A., Hicks, D.M., Kunin, W.E., Mitschunas, N., Morse, H., Osgathorpe, L.M., Potts, S.G., Robertson, K.M., Scott, A.V., Staniczenko, P.P.A., Stone, G.N., Vaughan, I.P. & Memmott, J. (2019) A systems approach reveals urban pollinator hotspots and conservation opportunities. Nature Ecology & Evolution, 3, 363–373.

Bełcik, M., Woźniak, B. & Skórka, P. (2025) Forest fragmentation and heterogeneity shape the occurrence of woodpecker species in Central Europe. Scientific Reports, 15, 21660.

Beninde, J., Veith, M. & Hochkirch, A. (2015) Biodiversity in cities needs space: a meta-analysis of factors determining intra-urban biodiversity variation. Ecology Letters, 18, 581–592.

Bovyn, R.A., Lordon, M.C., Grecco, A.E., Leeper, A.C. & LaMontagne, J.M. (2019) Tree cavity availability in urban cemeteries and city parks. Journal of Urban Ecology, 5.

Catalina-Allueva, P. & Martín, C.A. (2021) The role of woodpeckers (family: Picidae) as ecosystem engineers in urban parks: a case study in the city of Madrid (Spain). Urban Ecosystems, 24, 863–871.

Croeser, T., Sharma, R., Weisser, W.W. & Bekessy, S.A. (2024) Acute canopy deficits in global cities exposed by the 3-30-300 benchmark for urban nature. Nature Communications, 15, 9333.

Dennis, E.B., Morgan, B.J., Brereton, T.M., Roy, D.B. & Fox, R. (2017) Using citizen science butterfly counts to predict species population trends. Conservation Biology.

Fairbairn, A.J., Meyer, S.T., Mühlbauer, M., Jung, K., Apfelbeck, B., Berthon, K., Frank, A., Guthmann, L., Jokisch, J. & Kerler, K. (2024) Urban biodiversity is affected by human-designed features of public squares. Nature Cities, 1, 706–715.

Figarski, T. & Kajtoch, Ł. (2018) Hybrids and mixed pairs of Syrian and great-spotted woodpeckers in urban populations. Journal of ornithology, 159, 311–314.

Fontaine, B., Bergerot, B., Le Viol, I. & Julliard, R. (2016) Impact of urbanization and gardening practices on common butterfly communities in France. Ecology and Evolution, 6, 8174–8180.

Kajtoch, Ł. & Figarski, T. (2017) Comparative distribution of Syrian and great spotted woodpeckers in different landscapes of Poland. Folia Zoologica, 66, 29–36.

Korányi, D., Gallé, R., Donkó, B., Chamberlain, D.E. & Batáry, P. (2021) Urbanization does not affect green space bird species richness in a mid-sized city. Urban Ecosystems, 24, 789–800.

Kotaka, N. & Matsuoka, S. (2002) Secondary users of Great Spotted Woodpecker (Dendrocopos major) nest cavities in urban and suburban forests in Sapporo City, northern Japan. Ornithological Science, 1, 117–122.

Lepczyk, C.A., Aronson, M.F.J., Evans, K.L., Goddard, M.A., Lerman, S.B. & Macivor, J.S. (2017) Biodiversity in the City: Fundamental Questions for Understanding the Ecology of Urban Green Spaces for Biodiversity Conservation. Bioscience, pp. 799–807.

Melliger, R.L., Rusterholz, H.-P. & Baur, B. (2017) Habitat- and matrix-related differences in species diversity and trait richness of vascular plants, Orthoptera and Lepidoptera in an urban landscape. Urban Ecosystems, 20, 1095–1107.

Morrison, J.L. & Chapman, W.C. (2005) Can urban parks provide habitat for woodpeckers? Northeastern Naturalist, 12, 253–262.

Mühlbauer, M., Weisser, W.W., Müller, N. & Meyer, S.T. (2021) A green design of city squares increases abundance and diversity of birds. Basic and Applied Ecology, 56, 446–459.

Pasinelli, G. (2006) Population biology of European woodpecker species: a review. Annales Zoologici Fennici, 43, 96–111.

Rega-Brodsky, C.C., Aronson, M.F.J., Piana, M.R., Carpenter, E.-S., Hahs, A.K., Herrera-Montes, A., Knapp, S., Kotze, D.J., Lepczyk, C.A., Moretti, M., Salisbury, A.B., Williams, N.S.G., Jung, K., Katti, M., MacGregor-Fors, I., MacIvor, J.S., La Sorte, F.A., Sheel, V., Threfall, C.G. & Nilon, C.H. (2022) Urban biodiversity: State of the science and future directions. Urban Ecosystems.

Sandström, U.G., Angelstam, P. & Khakee, A. (2006) Urban comprehensive planning–identifying barriers for the maintenance of functional habitat networks. Landscape and urban planning, 75, 43–57.

Sandström, U.G., Angelstam, P. & Mikusiński, G. (2006) Ecological diversity of birds in relation to the structure of urban green space. Landscape and Urban Planning, 77, 39–53.

Schütz, C., Reckendorfer, W. & Schulze, C.H. (2017) Local quality versus regional connectivity—habitat requirements of wintering woodpeckers in urban green spaces. Journal of Urban Ecology, 3.

Stański, T., Czeszczewik, D., Stańska, M. & Walankiewicz, W. (2020) Foraging behaviour of the Great Spotted Woodpecker Dendrocopos major in relation to sex in primeval stands of the Białowieża National Park. Acta Ornithologica, 55, 120–128.

Sweet, F.S.T., Apfelbeck, B., Hanusch, M., Garland Monteagudo, C. & Weisser, W.W. (2022) Data from public and governmental databases show that a large proportion of the regional animal species pool occur in cities in Germany. Journal of Urban Ecology, 8, juac002.

Sweet, F.S.T., Noack, P., Hauck, T.E. & Weisser, W.W. (2023) The Relationship between Knowing and Liking for 91 Urban Animal Species among Students. Animals, 13.

Venables, W.N. & Ripley, B.D. (2002) Modern Applied Statistics with S. Fourth Edition. Springer, New York.

von Blotzheim, U.N.G., Bauer, K. & Bezzel, E. (1966) Handbuch der Vögel Mitteleuropas. Akademische Verlagsgesellschaft.

Wagenführ, A. & Scholz, F. (2018) Taschenbuch der Holztechnik. Carl Hanser Verlag GmbH Co KG.

Weisser, W.W. & Hauck, T.E. (2024) Animal-Aided Design–planning for biodiversity in the built environment by embedding a species’ life-cycle into landscape architectural and urban design processes. Landscape Research, 1–22.

Winkler, H. & Short, L.L. (1978) A comparative analysis of acoustical signals in pied woodpeckers (Aves, Picoides). Bulletin of the AMNH; v. 160, article 1.

Zanne, A.E., Lopez-Gonzalez, G., Coomes, D., Ilic, J., Jansen, S., Lewis, S., Miller, R., Swenson, N., Wiemann, M. & Chave, J. (2009) Global wood density database. Dryad.

## References

Fairbairn, A.J., Meyer, S.T., Mühlbauer, M., Jung, K., Apfelbeck, B., Berthon, K., Frank, A., Guthmann, L., Jokisch, J., Kerler, K., Müller, N., Obster, C., Unterbichler, M., Webersberger, J., Matejka, J., Depner, P. & Weisser, W.W. (2024) Urban biodiversity is affected by human-designed features of public squares. Nature Cities, 1, 706–715.

Mühlbauer, M., Weisser, W.W., Apfelbeck, B., Müller, N. & Meyer, S.T. (2024) Bird guilds need different features on city squares. Oecologia, 82, 23–35.

